# Susceptibility of European freshwater fish to climate change: species profiling based on life-history and environmental characteristics

**DOI:** 10.1101/355875

**Authors:** Ivan Jarić, Robert J. Lennox, Gregor Kalinkat, Gorčin Cvijanović, Johannes Radinger

## Abstract

Climate change is expected to strongly affect freshwater fish communities. Combined with other anthropogenic impacts, the impacts will alter species distributions and contribute to population declines and local extinctions. To provide timely management and conservation of fishes, it is relevant to identify species that will be most impacted by climate change and those that will be resilient. Species traits are considered a promising source of information on characteristics that influence resilience to various environmental conditions and impacts. We collated life history traits and climatic niches of 443 European freshwater fish species and compared those identified as susceptible to climate change to those that are considered to be resilient. Significant differences were observed between the two groups in their distribution, life-history and climatic niches, with climate-change susceptible species being distributed more southwardly within Europe, and being characterized by higher threat levels, lower commercial relevance, lower vulnerability to fishing, smaller body size and warmer thermal envelopes. We establish a list of species revealed to be of highest priority for further research and monitoring regarding climate change susceptibility within Europe. The presented approach represents a promising tool, to quickly assess large groups of species regarding their susceptibility to climate change and other threats, and to identify research and management priorities.

## Introduction

As ectothermic organisms, fishes are intimately linked to local climatic conditions through physiological mechanisms that delimit tolerance or resilience (Comte & Olden, 2017a). Zoogeography of fishes is therefore greatly influenced by the average and spread of temperatures experienced in a given watershed (Pörtner & Farrell, 2008; Isaak & Rieman, 2013). Relative to seas and oceans, freshwater habitats are more drastically impacted by changes in climate, especially due to changes in temperature and flow, and climate change is projected to strongly affect freshwater fish communities (O’Reilly et al., 2003; Buisson et al., 2008; Graham & Harrod, 2009; Harrod, 2016; Radinger et al., 2017). Combined with other anthropogenic impacts (e.g. land use change and thermal pollution; Radinger et al., 2016; Raptis et al., 2017), climate change will restrict or redraw thermal envelopes, contribute to population declines and local extinctions, and overall shifts in the distribution of species. Riverine fish species losses due to climate change and reduced water discharge are predicted to reach 75% in some river basins (Xenopoulos et al., 2005). Phenological changes in fish behaviour (Otero et al., 2014; Dempson et al., 2017; Hovel et al., 2017) have been also detected and emphasize the powerful changes imposed by a changing climate. In Europe, there is a broad range of climatic conditions experienced across the landscape and a diverse ichthyofauna distributed throughout the lakes and rivers (Ficke et al., 2007). Within the IUCN (International Union for Conservation of Nature and Natural Resources) Red List, as much as 33% of European freshwater fish species are recognized as threatened by climate change (IUCN, 2017).

Efforts to preserve ecosystem integrity must focus on maintaining species richness and diversity to ensure that the services provided by freshwater ecosystems are maintained. Conservation is often limited by funding and therefore must undergo triage to identify priorities and allocate resources efficiently (McDonald-Madden et al., 2011). To provide timely management and conservation and allocate resources efficiently, it is important to identify those species that will be most impacted by climate change and those that might be rather resilient. Species traits are considered as a promising source of information on characteristics that influence resilience to various environmental conditions and impacts (Jiguet et al., 2007; Comte & Olden, 2017b). Species traits represent any morphological, physiological or phenological feature that is measurable at the individual level of a species (Floeter et al., 2018). Trait-based evaluation has been demonstrated to be linked to the risk status of species and can be used to investigate mechanisms that contribute to imperilment, make predictions about unassessed species, or rank and prioritize species based on their relative risk (Olden et al., 2007; Bland & Böhm, 2016; Comte & Olden, 2018).

Here we assess various ecological and life-history characteristics of European freshwater fish species to identify traits that are characteristic for those that are susceptible to the effects of climate change. Automated scraping of an online trait database and calculation of climate envelopes using IUCN range maps overlaid on climate maps allowed us to collate species-specific data on life history, distribution, climatic niches, as well as data on threat and economic status. This allowed us to compare species identified as susceptible to climate change with those that are considered to be resilient. Results of the study will contribute to a better understanding of the expected climate change effects on European freshwater fish fauna. We also establish a list of European species of highest priority for further research and monitoring regarding climate change susceptibility. The method allows to extrapolate results and characterize rare and less studied species, with scarce autecological information.

## Materials and methods

### Dataset

Our analysis comprised comparisons of in total 443 European freshwater fishes between those that were identified as threatened by climate change (n=148) within the IUCN Red List Database (IUCN, 2017) and those without climate change listed as a threat (n=295).

A list of native European freshwater fish species belonging to 25 families, mainly to Cyprinidae (45%) and Salmonidae (20%), was obtained from the IUCN Red List database (IUCN, 2017). It included both exclusively freshwater species, as well as those that partly enter brackish and saltwater. Obtained data also comprised IUCN Red List classification and maps of their distributional range within Europe. In addition, we obtained information whether climate change was indicated as one of the threats for each species, which is based on threat analyses and expert judgement by IUCN species experts. Overall, the dataset comprised 33% species which were categorized as susceptible to climate change.

In addition, for each species we collated trait information related to their life history, ecology, fishery and threat status, and spatial and bioclimatic data variables (Table 1). Life-history data were obtained from the FishBase database (Froese & Pauly, 2017) by using the rfishbase R package (Boettiger et al., 2012, 2017). Traits with low data coverage (i.e. those that were available for less than 1% of all species) were excluded from the analysis. Bioclimatic spatial data were obtained from the MERRAclim database (Vega et al., 2017) and included 19 variables related to temperature and humidity (Table 1). Global Human Footprint map (map of anthropogenic impacts on the environment) was obtained from WCS and CIESIN (2005) and the spatial elevation data were obtained from USGS (2010). Based on the distributional range maps of each species (IUCN, 2017), mean values within each species’ range were estimated for each of the spatial variables using the intersect tool in ArcGIS (version 10.5) and the *extract*function in the R (version 3.4.3; R Core Team, 2017) package raster (version 2.6-7; Hijmans, 2017). Range maps were also used to estimate the number of watersheds covered by each species based on WRI (2006), as well as the area and coordinates of the range centroid for each species. General descriptive statistics and information on data sources of all variables used in the analysis is presented in Table 1 and Supplementary material S1. The complete dataset is presented in Supplementary material S2.

**Table 1.**
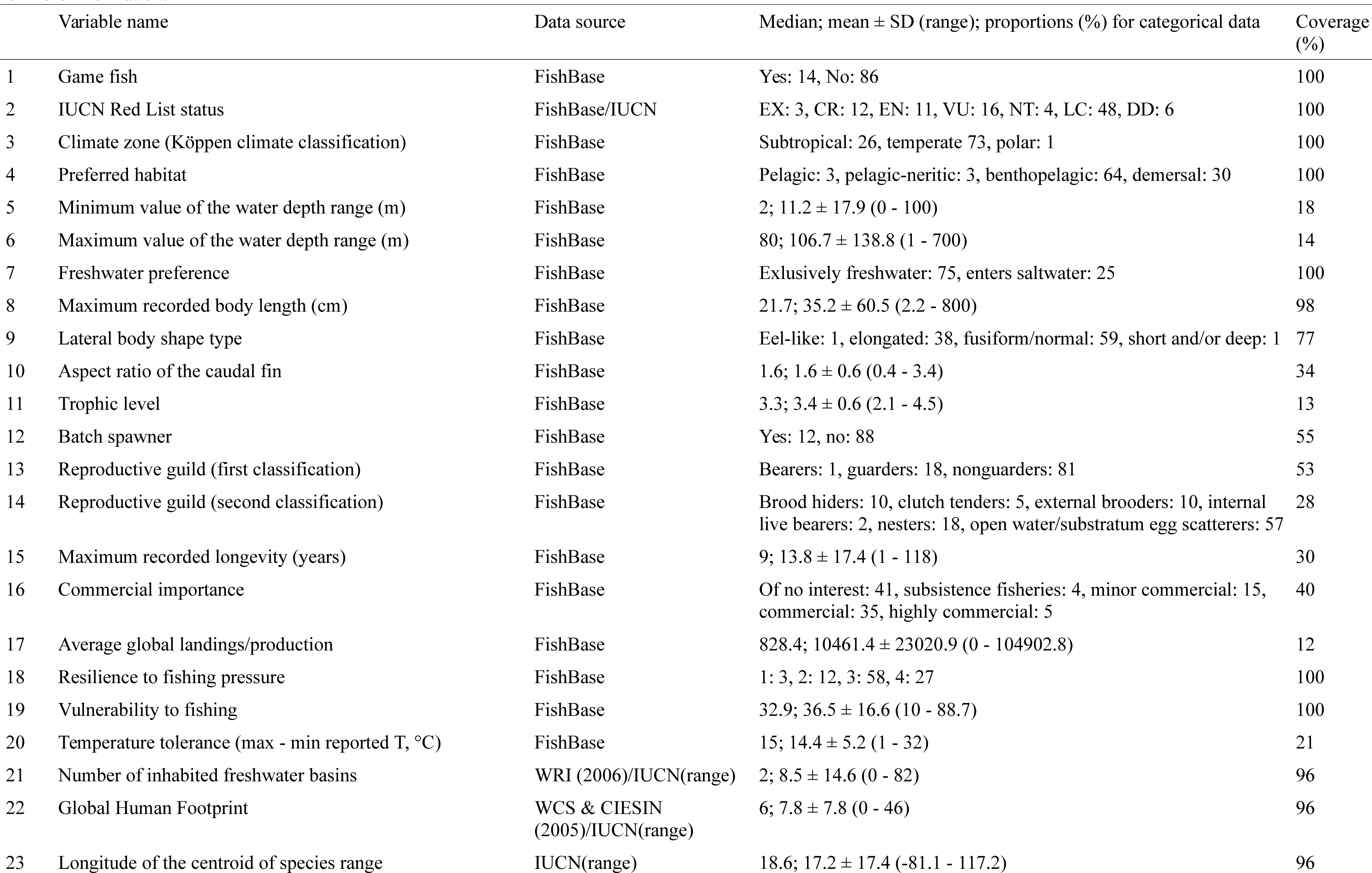
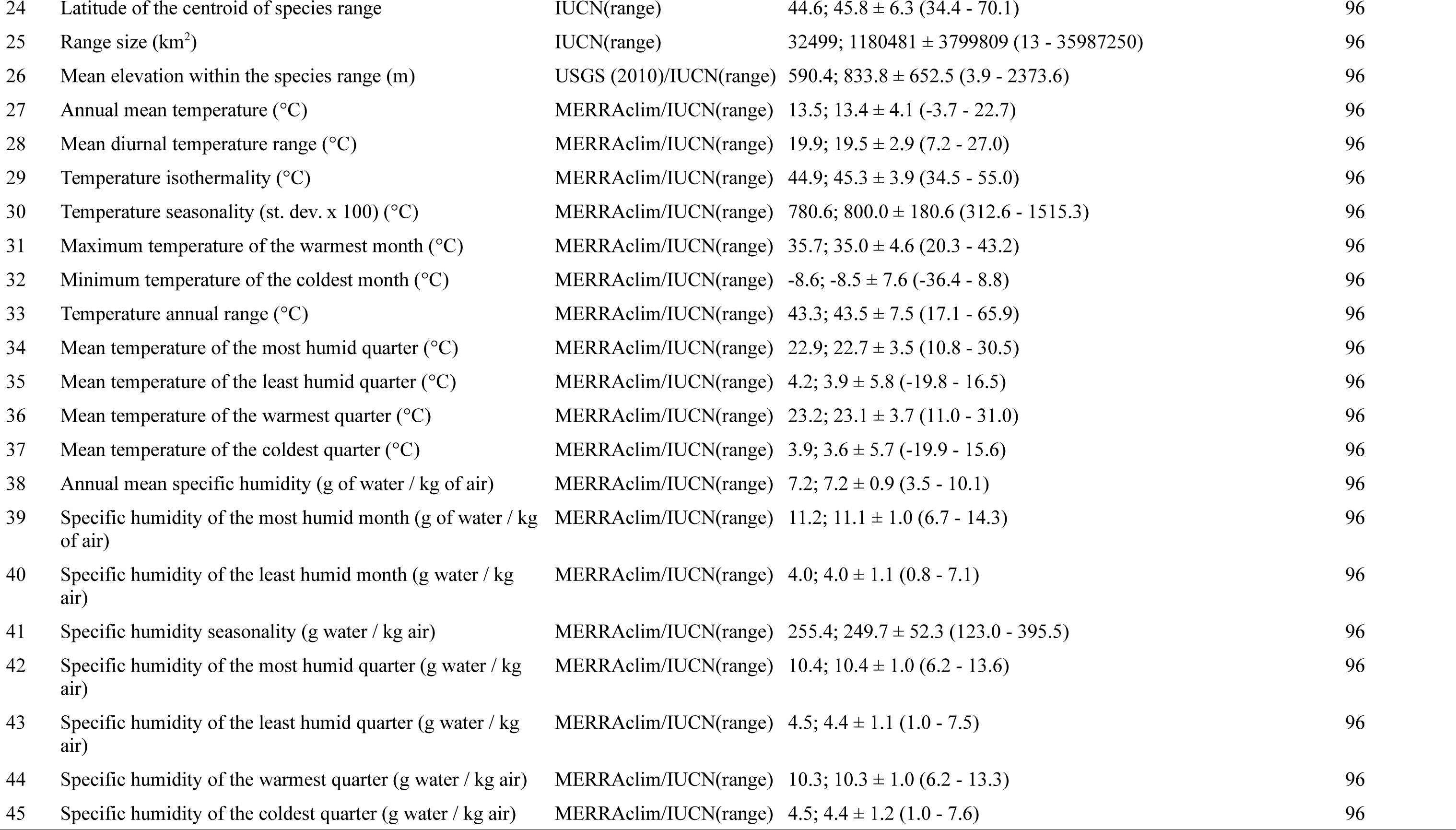
Variables used in the analysis, with their data sources, general descriptive statistics and coverage (proportion of species with available data). See Supplemetary material I for more information.

### Statistical Analysis

We calculated boosted regression trees (BRT) to evaluate the relationship between the membership of a species to the group of susceptible vs. non-susceptible species and the 45 explanatory variables. BRT are a statistical learning method that combines and averages (boosting) many simple single regression trees to form collective models of improved predictive performance (Elith et al., 2008). BRT can accommodate continuous and categorical variables, are not affected by missing values or transformation or outliers and are considered to effectively select relevant variables, identify variable interactions and avoid overfitting (Elith et al., 2008, Radinger et al., 2015).

Specifically, we first fitted an initial global BRT model (R package *dismo*, *gbm.step*, version 1.1-4; Hijmans et al., 2017) using the complete set of explanatory variables. An automatized stepwise backward selection of explanatory variables (*gbm.simplify*) was applied to eliminate non-informative variables based on model-internal cross-validation of changes in a models’ predictive deviance (Hijmans et al., 2017). Thereafter, we calculated a final BRT model (*gbm.step*) based on the selected set of explanatory variables. For all BRT modeling steps, tree complexity and learning rate was set to 3 and 0.001, respectively, to achieve the recommended number of more than 1000 regression trees (Elith et al., 2008). All other model settings were set to default or were automatically adjusted by the boosting algorithm. We calculated a 10-fold cross validation of the BRT model as already implemented in the algorithm. In addition, we extracted the mean AUC (area under the receiver operating characteristic curve) as a measure of the model’s predictive quality. The AUC is a threshold-independent rank-correlation coefficient with high values typically indicating a strong agreement between the model prediction and the membership of species to the susceptible vs. non-susceptible group (Hijmans & Elith, 2017).

The relative importance (%) of each explanatory variable in the final BRT model was quantified based on the number of times each variable was used for splitting, weighted by the squared improvement at each split and averaged over all trees (Elith et al., 2008). For BRT models with Gaussian distribution the relative variable importance equals the reduction of squared error attributable to a given variable. Differences between groups were also assessed by bootstrapping, by sampling each group independently and estimating the difference based on confidence intervals (functions *two.boot*and *boot.ci*, R package *simpleboot*, version 1.1-3; Peng 2008). Differences were considered to be significant if 95% confidence intervals did not overlap with zero.

Subsequently, species were ranked based on the subset of variables selected by the BRT analysis (i.e., those with >1% variable importance score), and weighed by the importance of each variable, in order to estimate their position along the climate change susceptibility continuum. For each species, the value of each variable was standardized based on its position between the minimum (*t*_min_) and maximum values (*t*_max_) observed in the dataset, with 0-1 possible range, and multiplied by the importance score (*I _x_*) of the given variable:

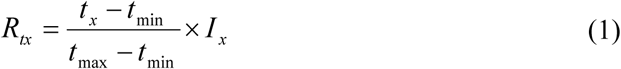

where *R _tx_* represents the rank value of variable *t*in species *x*, and *t _x_* is the value of variable *t*for species *x*. For variables where the lower endpoint (*t*_min_) is associated with the climate change susceptibility, equation should be adjusted as follows:

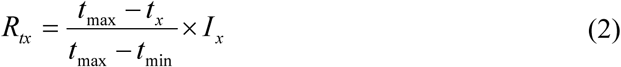

Summing of all ranking scores across all variables yielded the total species ranking score, which can range from 0 to 100, with higher values indicating stronger climate change susceptibility.

## Results

Our analyses indicated substantial differences between the two groups, climate change susceptible versus non-susceptible species. The BRT analysis selected 35 most relevant variables, which were subsequently assessed for their relative importance to discriminate between the two groups (Figure 1 and Supplementary material S3). The BRT model with the selected set of explanatory variables was successfully modeled (Supplementary material S3) with a good cross-validated AUC of 0.87 (standard error = 0.014). Out of all explanatory variables, latitude of the species range centroid was selected as by far the most relevant variable (31% variable importance), followed by the IUCN Red List classification (8%), commercial relevance (6%) and vulnerability to fishing (6%). Climate susceptible species were characterized by more southwardly positioned distribution range centroids (41.6º vs. 47.8º N as a mean value in susceptible and non-susceptible species, respectively), smaller range sizes (175 x 10^3^km^2^vs. 1686 × 10^3^km^2^), and lower elevations within their ranges (717.7 m vs. 892.2 m a.s.l.), with a higher proportion of exclusively freshwater species (93% vs. 66%; Figure 2). Susceptible species were also characterized by a smaller maximum body length (23.4 cm vs. 41.0 cm), higher proportion of threatened species (63% vs. 31%), lower proportion of commercially relevant species (25% vs. 74% of highly commercial, commercial and minor commercial species), and lower vulnerability to overfishing (32.6 vs. 38.5 vulnerability index; Figure 3), as well as by higher temperature-related values (Supplementary material S4). Bootstrapping indicated significant differences between the groups in each of the variables. Species that are susceptible to climate change are mainly distributed within the Mediterranean region, while the non-susceptible species distribution mainly covers central and northern European regions, as well as the Carpathian region (Figure 4).

**Figure 1.**
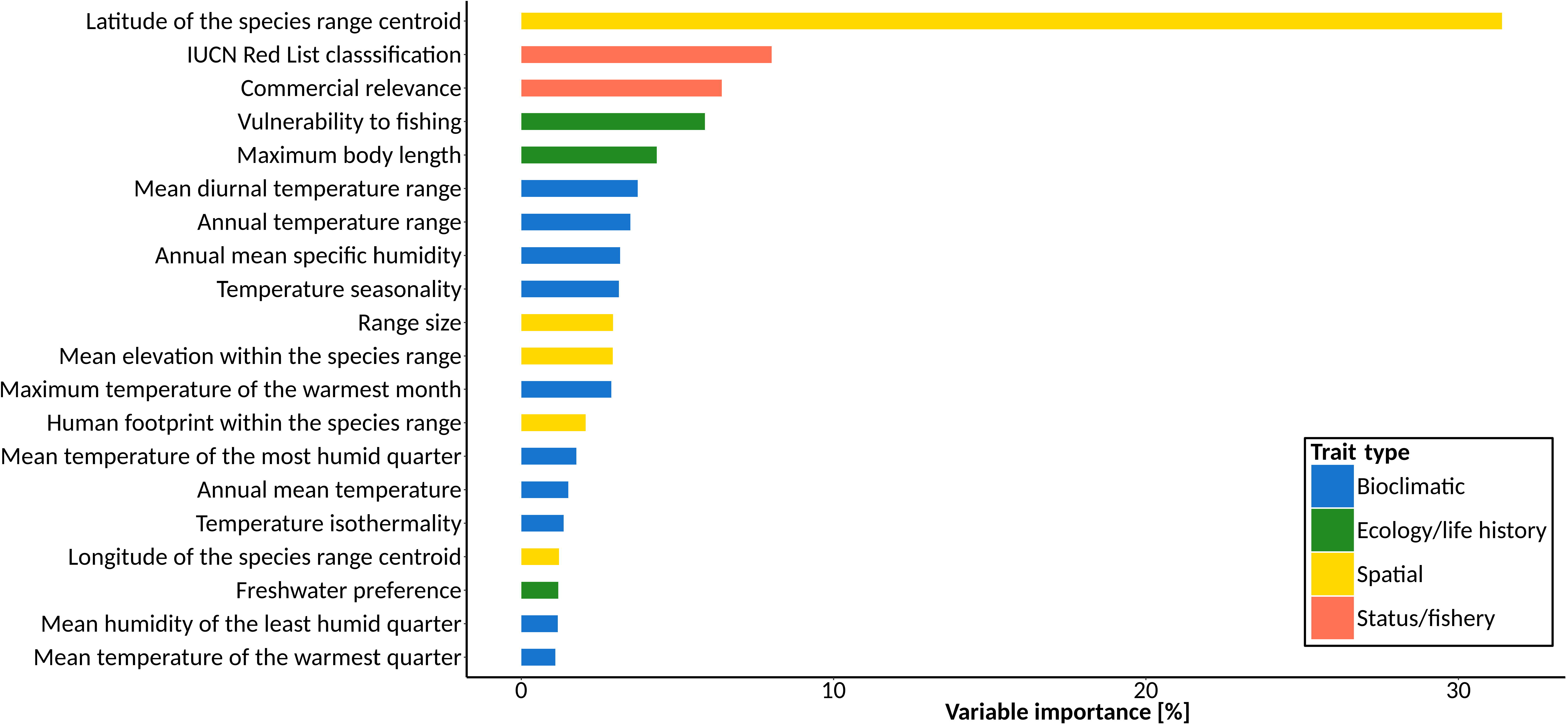
Variables selected by the boosted regression tree (BRT) model as the most relevant descriptors of climate change susceptibility in European freshwater fish species; 20 most relevant variables are presented, which together account for 90% of the total relative variable influence.

Species ranking based on the association of their traits with climate change susceptibility characteristics is presented in Table 2 and Supplementary material S5. The five top-ranked climate susceptible species were Acheron spring goby (*Knipowitschia milleri*), Corfu toothcarp (*V a l enc i a letourneuxi*), *Iberochondrostoma almacai*, Evia barbel (*Barbus euboicus*) and Malaga chub (*Squalius malacitanus*). Most of the species with the high climate-susceptibility ranks are also classified as highly threatened according to the IUCN classification (Table 2). Interestingly, the highest ranked species, *K. milleri*, was not classified within the IUCN Red List as threatened by climate change. Other high-ranking species that were not recognized as threatened by climate change were Malaga chub (*Squalius malacitanus*), Almiri toothcarp (*Aphanius almiriensis*), and Trichonis dwarf goby (*Economidichthys trichonis*). Species with the lowest ranking scores, i.e. with low climate change susceptibility, were humpback whitefish (*Coregonus pidschian*), Arctic flounder (*Liopsetta glacialis*), northern pike (*Esox lucius*), burbot (*Lota lota*), and European perch (*Perca fluviatilis*). A complete list of all species’ rankings is presented in Supplementary material S5.

**Table 2.**
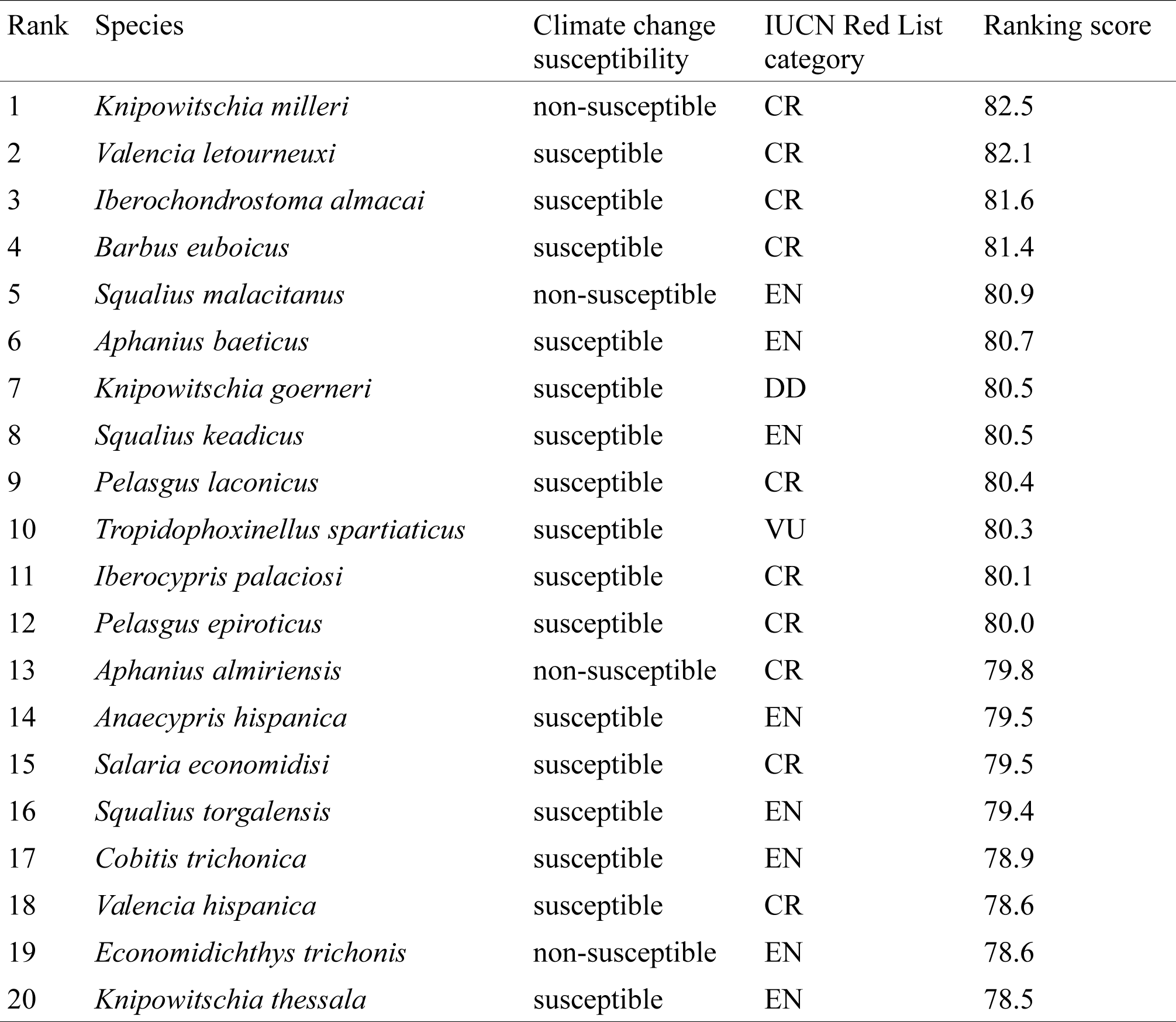
European freshwater fish species with the highest ranking scores, estimated based on the association of their traits with the climate change susceptibility characteristics, as indicated by the BRT model. Complete ranking list of all species is presented in Supplementary material 5.

## Discussion

In the present study, significant differences in life-history and climatic niches were observed between the European freshwater species susceptible to climate change and those that are not, such as species body size, range size, distribution and thermal envelopes. The latitude of the species range centroid was by far the most influential trait. Overall, southern regions with the warmer, Mediterranean climate comprised a higher proportion of species susceptible to climate change (Figure 4). These results support recent findings that the species from lower latitudes and tropical, warm-water habitats, are at greater risk from climate change and warming (Payne & Smith, 2016; Payne et al., 2016; Comte & Olden, 2017b). In such species, evolved towards higher upper thermal tolerances, specialization to thermal extremes is accompanied by a reduced physiological flexibility and adaptation capacity to respond to changing environmental conditions (Payne & Smith, 2016; Payne et al., 2016; Comte & Olden, 2017b). Such heat-tolerant species are also adapted to temperatures close to their physiological thermal limits, with a narrow safety margin for further temperature increases (Sinclair et al., 2016; Comte & Olden, 2017b). Freshwater basins in Southern Europe are also of particular conservation concern due to an elevated pressure by a range of anthropogenic impacts that further exacerbate effects of climate change, such as pollution, water resource development and consumption, and biological invasions (Xenopoulos et al., 2005; Clavero & García-Berthou, 2005; Walther et al., 2009; Vörösmarty et al., 2010; Comte & Olden, 2017a).

Climate-susceptible species were also characterized by a smaller body and range size (Figures 2, 3). These traits, which are also related to a lower dispersal ability (Radinger & Wolter, 2014), are well recognized as predictors of climate change susceptibility in fish (e.g. Ficke et al., 2007; Isaak & Rieman, 2013; Chessman, 2013; Pearson et al., 2014; Radinger et al., 2017). Smaller-bodied fish are facing elevated overall extinction risk in freshwater habitat due to multiple threats, such as habitat loss and fragmentation (Olden et al., 2007; Kalinkat et al., 2017; Kopf et al., 2017), which explains higher threat level observed in climate-susceptible species in the present study. Observed lower commercial relevance and lower vulnerability to fishing of climate-susceptible species both stem from a lower body size and related faster life history of such species.

**Figure 2.**
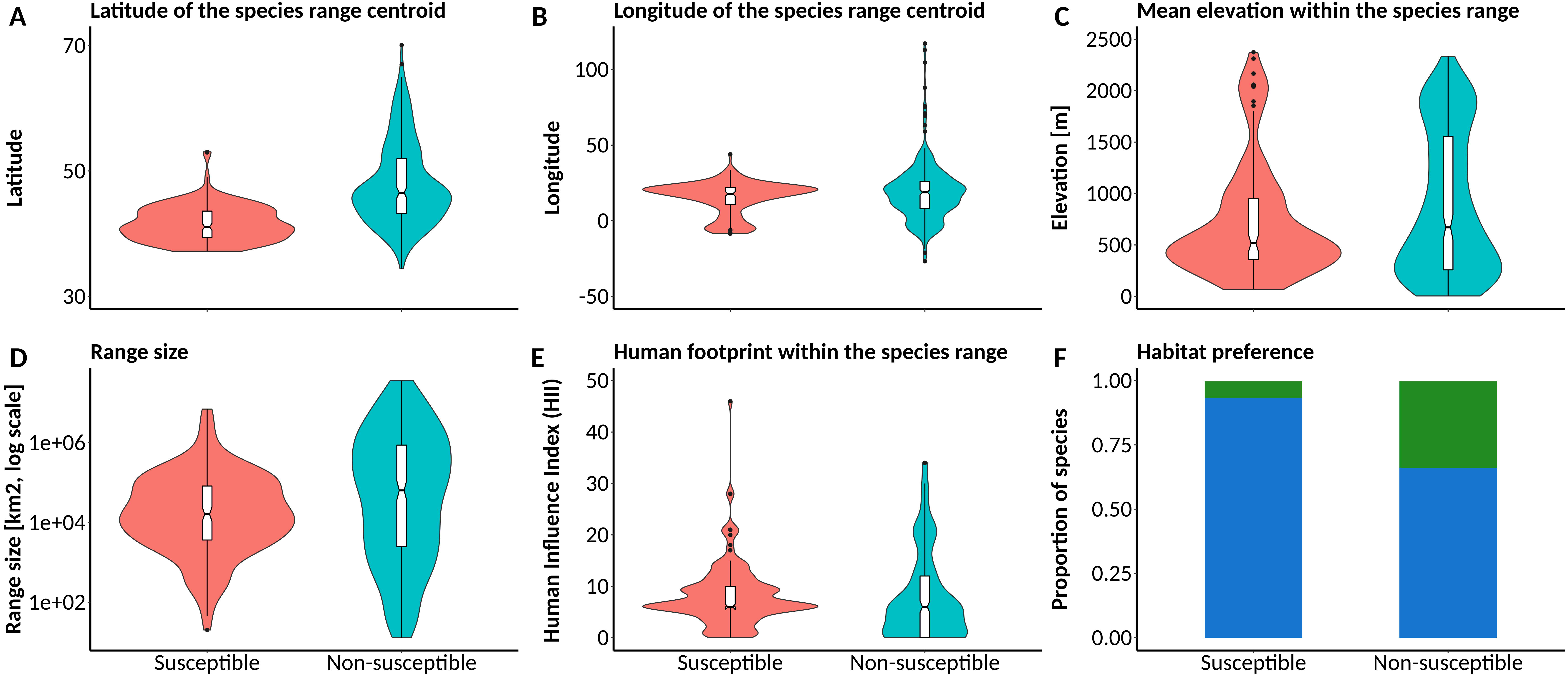
Violin-boxplots and barplots of the most relevant spatial variables in European freshwater fish species indicated as either susceptible (n = 148) or non-susceptible (n = 295) to climate change. Habitat preference: blue - exclusively freshwater species, green – species that also enter saltwater.

**Figure 3.**
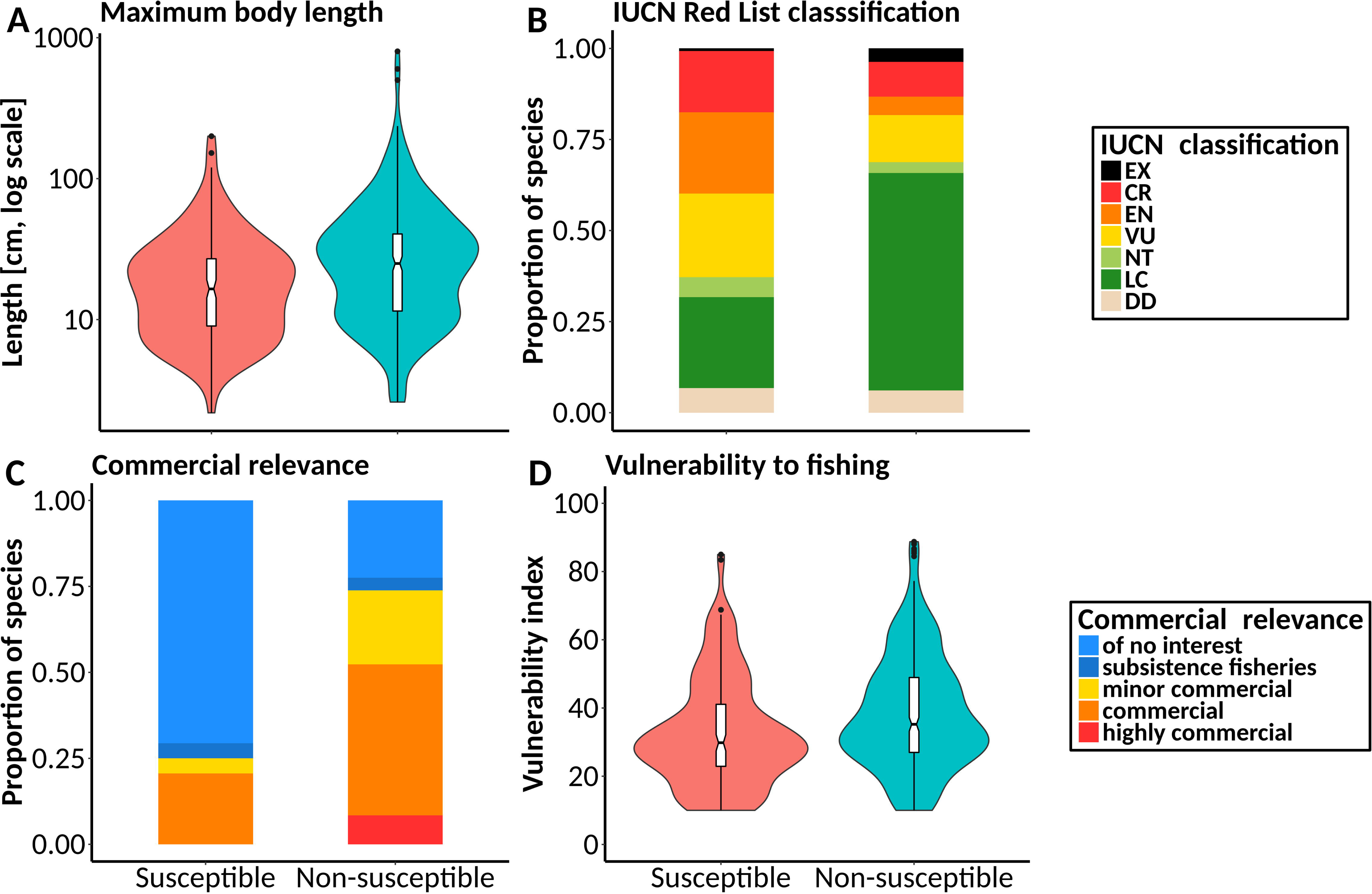
Violin-boxplots and barplots of the most relevant life history traits and variables related to threat and commercial status in European freshwater fish species indicated as either susceptible (n = 148) or non-susceptible (n = 295) to climate change.

**Figure 4.**
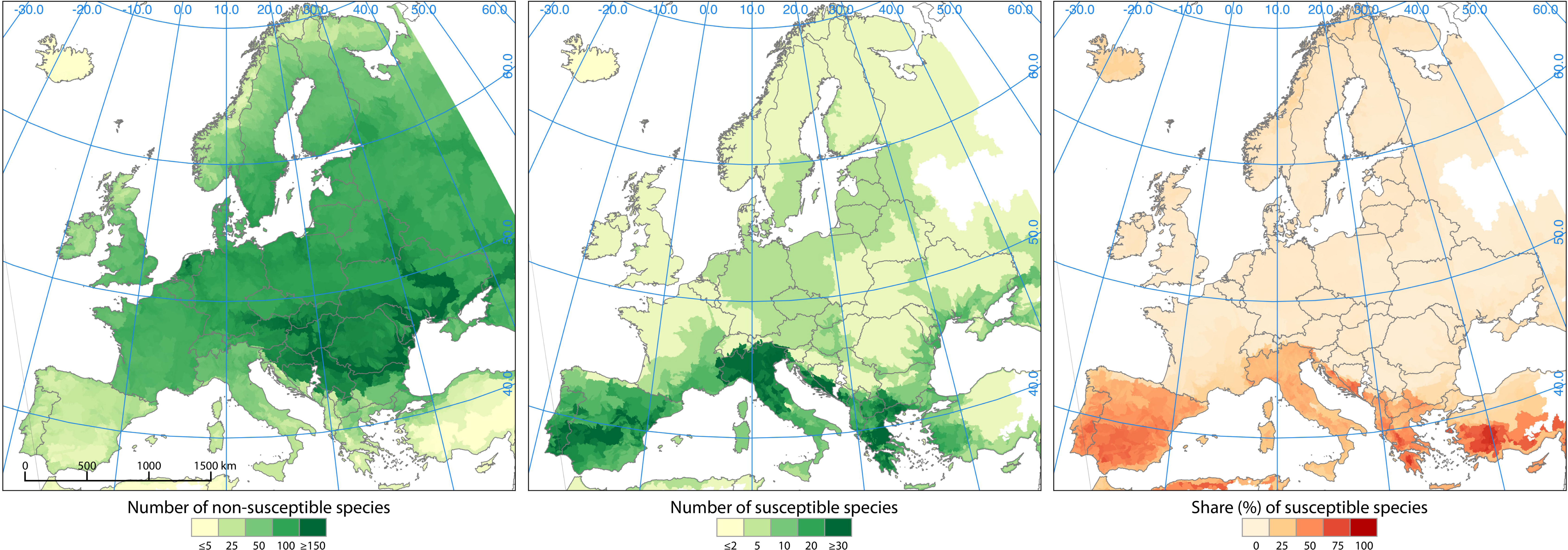
Richness of freshwater fish species across Europe indicated as either susceptible (middle panel) or non-susceptible (left panel) to climate change, and the relative share of susceptible species in the local total species richness (right panel).

It is important to acknowledge certain limitations of the data sources used in this study, such as species and trait coverage, reliability of methods applied for threat and extinction risk classification, and potential assessors’ biases (Clavero & García-Berthou, 2005; Keith et al., 2014; Trull et al., 2018). Furthermore, species that are not classified within IUCN Red List as threatened by climate change can comprise also those that are not yet assessed for their major threats. Nevertheless, the focus of our study on a well-studied group such as European species ensured that the basic life history data and IUCN Red List assessments were mostly available (Kopf et al., 2017). IUCN Red List is sometimes considered to understate or improperly account for climate change as a threat, mostly due to ambiguous definitions and criteria (Trull et al., 2018). However, recent studies have indicated that the IUCN classification is more efficient in detecting species vulnerable to climate change than anticipated (Keith et al., 2014; Pearson et al., 2014). Notwithstanding all the caveats, the databases used in the present study still represent the most comprehensive sources of data and the best available knowledge (Olden et al., 2007; Vega et al., 2017).

Trait-based risk assessments are increasingly used for species profiling (Pacifici et al., 2015; Liu et al., 2017; MacLean & Beissinger, 2017). The approach presented in this study might be considered a valid and promising approach to be used as a screening tool, i.e. to quickly assess large groups of species regarding their susceptibility to climate change and other threats based on species traits, and to identify research and management priorities. Our results indicate that the European environmental policy related to climate change mitigation and adaptation (EEA, 2012, 2017) should be mainly focused on Mediterranean region. This is especially important since this region is also predicted to have the highest frequency of droughts and extreme high temperatures, strongest reduction in precipitation and river discharges, the highest aggregate climate change impact and the lowest adaptation capacity (Milly et al., 2005; Dankers & Feyen, 2008; Fischer & Schär, 2010; ESPON Climate, 2011; Stagge et al., 2011; Rojas et al., 2012; Jacob et al., 2014; Russo et al., 2014). Moreover, Mediterranean region was also identified as a European priority area regarding climate change impacts for other species groups. A similar distributional pattern of species susceptible to climate change was previously also reported for aquatic insects such as Plecoptera, Ephemeroptera and Trichoptera (Hering et al., 2009; Conti et al., 2014), mammals (Levinsky et al., 2007), as well as for terrestrial species in general (Pacifici et al., 2015).

Species ranking conducted here indicated priority species for further research and monitoring regarding climate change (e.g. *V. letourneuxi*, *I. almacai*and *B. euboicus*; Table 2). Moreover, it also identified species whose IUCN Red List status potentially needs to be reconsidered or updated, such as highly ranked but apparently non-susceptible species (e.g. *K. milleri*), or highly ranked species without a proper threat category (e.g. *K. goerneri*, classified as Data Deficient species). As such, it has a potential to be used as a “Robin Hood Approach” (Punt et al., 2011), where assessments based on information-rich species are used to evaluate and categorize those that are information-poor. There is a need for climate change vulnerability assessments that would be based on quantitative approaches and consistent set of criteria, such as trait-based approaches advocated by IUCN (Foden et al., 2013; Trull et al., 2018). The approach presented here could be a good step in that direction.

## Acknowledgements

IJ acknowledges the sponsorship provided by the The J. E. Purkyně Fellowship of the Academy of Sciences of the Czech Republic and funding from the European Union’s Horizon 2020 research and innovation programme through the project ClimeFish (grant No. 677039), as well as funding provided by the Alexander von Humboldt Foundation and the German Federal Ministry of Education and Research (BMBF). GK acknowledges funding from the BMBF through the project “GLANCE” (Global Change Effects in River Ecosystems; 01 LN1320A). JR acknowledges funding through the 2015-2016 BiodivERsA COFUND call for research proposals and the Spanish Ministry of Economy, Industry and Competitiveness (project ODYSSEUS, BiodivERsA3-2015-26, PCIN-2016-168). GC acknowledges funding from the Ministry of Education, Science and Technological Development, Republic of Serbia (grant number 173045). Thanks are extended to Ryder Burt for assistance with geographic information processing.

